# Correlated responses to experimental evolution of increased post-infection survival in *Drosophila melanogaster*: Life-history trade-offs and reaction to novel stressors

**DOI:** 10.1101/2022.06.25.497416

**Authors:** Aparajita Singh, Aabeer Basu, Tejashwini Hegde, Biswajit Shit, Nitin Bansal, Ankita Chauhan, Nagaraj Guru Prasad

**Author notes:** These authors contributed equally to this study (Biswajit Shit, Nitin Bansal, Ankita Chauhan).

## Abstract

Evolution of increased immune defence is often limited by costs: correlated changes in other traits (viz. life-history traits) that otherwise reduce the fitness of the host organisms. Experimental evolution studies are useful for understanding the evolution of immune function, and correlated changes in other traits. We experimentally evolved replicate *Drosophila melanogaster* populations to better survive infection challenge with an entomopathogenic bacteria, *Enterococcus faecalis*. Within 35 generations of forward selection, selected populations showed marked increase in post-infection survival compared to ancestrally paired controls. We next measured various life-history traits of these populations. Our results show that the selected populations do not differ from control populations for larval development time and body weight at eclosion. No difference is also observed in case of fecundity and longevity (following the acute phase of infection), either when the flies are subjected to infection or when the flies were uninfected; although infected flies from all populations die much earlier compared to uninfected flies. Selected flies and control flies are also equally good at surviving abiotic stressors (starvation and desiccation), although infected flies from all populations are more susceptible to stress than uninfected flies. Therefore, we conclude that (a) *D. melanogaster* populations can rapidly evolve to be more immune to infection with *E. faecalis*; (b) evolution of increased defence against *E. faecalis* entails no life-history cost for the hosts; and (c) evolving defence against a biotic threat (pathogen) does not make flies more resistant to abiotic stressors.

## 1. INTRODUCTION

One of the central tenets of eco-immunology is that immune defense comes at a cost to the host (Sheldon and Verhulst 1996, Rolf and Siva-Jothy 2003, Schulenberg et al 2009). Previous authors have classified costs associated with immune function in diverse ways (viz., Schmid-Hempel 2003, McKean et al 2008, McKean and Lazzaro 2011, etc.). The cost of immune defense is expected to manifest in form of trade-offs with other organismal functions, such as reproductive output, life-history traits, and resistance to stressors (Lochmiller and Deerenberg 2000, Schmid-Hempel 2005).

We experimentally evolved replicate *Drosophila melanogaster* populations, selecting flies every generation for increased survival after being infected with a Gram-positive bacterial pathogen, *Enterococcus faecalis* (Singh et al 2021). Using these populations (and their ancestrally paired control populations; see Methods for details of the selection design), we explored if evolving increased defense against a bacterial pathogen leads to trade-offs with life-history traits in the hosts, by comparing flies from selected and control populations. We further explored if mounting a defense against the same bacterial pathogen is costly, by comparing infected and uninfected flies from the same population.

To test for trade-offs, we selectively measured life-history traits that have been previously demonstrated to have fitness consequences in *D. melanogaster* (reviewed Prasad and Joshi 2003). We measured larval development time and viability, and adult body weight, fecundity, and longevity. The adult traits (except body weight) were measured for both infected and uninfected flies from each population. In addition to these, we tested the response of evolved and control flies to novel biotic (intra-specific competition) and abiotic (starvation and desiccation) stressors. For the abiotic stressors too, we studied both infected and uninfected flies from each population.

Previous studies exploring cost of evolving increased immune defense in *D. malenogaster* have been equivocal on the matter. For example, both Kraaijeveld and Godfray (1997) and Fellowes et al (1998) demonstrated reduced larval competitive ability to be a cost of evolving defense against parasitoid infections. Similarly, flies evolved to better defend against the bacteria *Pseudomonas aeruginosa* have reduced egg viability and adult lifespan (Ye et al. 2009). On the other hand, no life-history costs were reported in two separate experimental evolution studies where flies were selected for increased defense against the bacteria *Pseudomonas entomophila* (Faria et al 2015, Gupta et al 2016). Flies experimentally co-evolved with *P. entomophila* also do not incur any life-history costs (Ahlawat et al 2022). There may be a few possible reasons for the inconsistency in results obtained in these studies, such as the pathogen/parasite used for selection, genetic architecture of the host population, specific life- history trait tested, and the amount of resource available to the host for allocation into different traits.

Studies exploring the cost of mounting an immune defense against pathogens and parasites have also similarly come up with discordant results. For example, trade-off between reproduction and immunity is a common expectation (Lawniczak et al 2007, Schwenke et al 2016), where infected hosts are expected to exhibit reduced reproductive effort (Lochmiller and Deerenberg 2000, Schmid-Hempel 2003, McKean et al 2008). But *D. melanogaster* females when infected with bacterial or viral pathogens are known to increase (Hudson et al 2020), decrease (Brandt and Schneider 2007, Linder and Promislow 2009, Gupta et al 2017), or show no change in their reproductive output (Kutzer and Armitage 2016, Kutzer et al 2018).

Our results show that evolution of increased defense against *E. faecalis*, in response to selection for increased post-infection survival, is not accompanied with life-history trade-offs. Selection history of the flies did not influence any of the life-history traits measured. There was a sex- specific effect of selection history on resistance to abiotic stressors: males of the selected populations were more resistant to both starvation and desiccation compared to males from the control population. Intra-specific competitive ability was affected by selection history of the host, but the difference between the selected and the control populations was determined by the intensity of competition.

## 2. MATERIALS AND METHODS

Life-history traits, and resistance to biotic and abiotic stressors, were measured for *Drosophila melanogaster* flies selected for improved post-infection survival against systemic infection with an entomopathogenic, Gram-positive bacteria, *Enterococcus faecalis*. The populations were named the *EPN populations*.

### 2.1. EPN populations: history and maintenance regime

The EPN populations was derived from ancestral BRB populations (for details refer Singh et al 2021). Briefly, from each replicate population of BRB_1-4_ three populations were derived: (i) E_1-4_, infected with *Enterococcus faecalis*; (ii) P_1-4_, pricking control; and (iii) N_1-4_, normal control. Populations having same numeral subscript shared a common recent ancestry and were treated as independent blocks (block 1, block 2 and so forth). Individual blocks were always handled together during selection and during experiments. Eggs were collected at a density of ∼70 eggs per vial (25mm diameter × 90 mm height) containing 6-8 ml of standard banana-jaggery-yeast food. Ten 10 such vials were collected for each population. These vials were reared under standard laboratory (12:12 light: dark cycle, 25 °C, 60% relative humidity) conditions until 12^th^ day post egg collection. By 10^th^-11^th^ day all flies eclose and had mated at least once by 12^th^ day (day of infection). Further handling depended on the type of population.

For E populations, on 12^th^ day post egg collection, every generation 200 females and 200 males were randomly picked out of total 700 flies. These flies were pricked on the dorsolateral surface of the thorax with Minutien pin (0.1mm Fine Scientific Tools, USA) dipped in bacterial suspension (refer section 2.2 Bacterial culture, for details). After infection flies were placed inside plexiglass cage (14 cm length x 16 cm width x 13 cm height) with food in 90 mm Petri plate. Fifty percent of the infected flies die within 96 hours of infection. Post 96 hours, fly cages were provided with oviposition food plates for 18 hours. Eggs were collected from these oviposition plates at a density of ∼70 eggs per vial (as mentioned above) to start next generation. This 18-hour window, 96 hours after infection, serves as the selection window for the EPN populations. The eggs produced by the surviving flies from each population during this window is used to set up the next generation.

Similarly, for P populations, 100 females and 100 males out of total 700 flies are pricked every generation with Minutien pin dipped in sterile 10 mM MgSO_4_. For N populations, 100 females and 100 males out of total 700 flies are sorted every generation under light CO_2_ anaesthesia and transferred to the plexiglass cage. Rest all the protocol are similar to E populations. There is negligible mortality in P and N populations. Post 96 hours, eggs are collected in similar way as for E populations to start the next generation.

### 2.2. Bacterial culture

The bacteria used in this study were *Enterococcus faecalis* (Ef, Gram-positive, grown at 37 °C, Lazzaro et al 2006). The bacterial stocks are maintained as 17% glycerol stocks frozen at -80 °C. An overnight primary culture of bacteria was set by inoculating a stab of bacterial glycerol stock in 10 ml lysogeny broth (Luria-Bertani-Miller, HiMedia) and incubating it at appropriate temperature with continuous mixing at 150 RPM (revolution per minute). Once this primary culture turned confluent (OD_600_ = 1.0-1.2), it was further diluted 100 times to set a secondary culture, and maintained at their respective conditions until it turned confluent again. This secondary culture was centrifuged, and bacterial pellets were resuspended in sterile 10 mM MgSO_4_ buffer to obtain desired OD_600_ for infection. This bacterial suspension was used to infect flies. Infection was done by dipping needle in the bacterial suspension or sterile 10mM MgSO_4_ buffer and pricking flies on the thorax.

For stock maintenance, E populations of EPN regime were infected with *E. faecalis*. Throughout the selection history of EPN, the pathogen infection dose was modulated to induce fifty percent mortality in E populations. This ensured a constant, directional selection process. Therefore, flies of zeroth generation of E were infected with *E. faecalis* at OD_600_=0.8 and when this experiment was done at generations 35-40 dose was increased to OD_600_=1.0.

### 2.3. Fly standardization

Experimental eggs were collected from flies that were cultured under common environmental conditions for one generation. This was done to account for any non-genetic parental effects (Rose 1984) and flies thus generated were called standardized flies.

Experimental eggs were collected from all three populations (E, P, and N) at a density of ∼70 eggs per vial and 10 such vials were established per population. The eggs were reared into the same development vials into adults, until day 12 PEL. These adults were then transferred to plexiglass cages (14x16x13 cm^3^) with food in Petri plates (60 mm diameter). Experimental eggs were collected from these population cages.

### 2.4. Post-infection survival and longevity

In this experiment, we measured the survival of flies from the EPN populations, under infected, sham-infected, and uninfected conditions, during the selection window (first 96 hours following infection), plus the remaining life-span of the flies that successfully survive the first 96 hours following treatment. This experiment was done after 35 generations of forward selection.

Standardized fly cages for each population (E_i_, Pi, and N_i_, where ‘i’ represents blocks 1-4) were provided with *ad libitum* yeast paste smeared on the top of the standard banana-jaggery-yeast food plate, three days prior to the egg collection. After two days, these yeasted food plates were replaced with oviposition plates for 18 hours. Eggs were collected from these oviposition plates at the density of approximately 70 eggs per vial, into 25 vials per population (E_i_, P_i_, and N_i_), with each vial containing 8 ml of banana-jaggery-yeast food. These *rearing* vials were incubated at standard maintenance conditions for next 12 days. Flies generally eclose by 10^th^- 11^th^ day post-egg-laying (PEL) in the rearing vials, and mate at least once by 12^th^ day PEL.

On 12^th^ day PEL, flies from each of the Ei, Pi, and Ni populations were randomly assigned to one of the three treatments:

a. *infected* treatment: 200 females and 200 males were randomly sampled from each population and were infected with *E. faecalis* at OD_600_ = 0.8 under light CO_2_ anaesthesia;
b. *sham-infected* treatment: 100 females and 100 males were randomly sampled from each population, and sham-infected with sterile needle dipped in sterile 10 mM MgSO_4_ solution; and,
c. *uninfected* treatment: 100 females and 100 males were just randomly sampled under light CO_2_ anaesthesia, and not subjected to any further manipulation.

After being subjected to different treatments, the flies were housed in plexiglass cages (14 cm × 16 cm × 13 cm) having *ad libitum* access to banana-jaggery-yeast food provided in Petri plates (60 mm diameter). Individual blocks were experimented upon on separate days.

Survivorship of the flies was monitored every 4-6 hours for the first 96 hours after infection, and after this period, mortality in the cages was recorded once a day until the last fly died in all cages. About fifty percent of the infected fly and almost all sham-infected and uninfected flies survived post 96-hours window. Fresh food plates were provided on alternate days. Altogether, 200 flies/sex/population/block were infected with *E. faecalis,* 100 flies/sex/population/block were sham-infected, and 100 flies/sex/population/block were maintained as uninfected controls.

### 2.5. Fecundity and hatchability

We measured the number of eggs produced by the females (fecundity) from the EPN populations, and what proportion of these eggs produced viable larva (hatchability), to test for the effect of selection history and infection status on fecundity and hatchability. The same experimental cages that were used for assaying post-infection survival was used here for measuring fecundity and hatchability.

Fecundity was assayed after 96 hours of infection treatment (identical to the time when eggs are collected for the next generation during maintenance of the selection regime). Each of the population cages (E_i_, P_i_, and N_i_ flies either infected, sham-infected, or uninfected) were provided with an oviposition plate (60 mm diameter), containing standard banana-jaggery-yeast food, for the female flies to lay eggs on for 18 hours. After 18 hours, these plates were withdrawn, labelled with the cage identity, and stored at -20°C for eggs to be counted later. Eggs on the surface of the food plates were counted visually using a light stereo microscope (Zeiss Stemi 2000) under 2.5X × 10X magnification. Per-female fecundity was calculated by dividing the number of eggs laid by the females during 18-hour window by the number of females alive in the respective cages at the start of the oviposition period.

Hatchability was assayed after 114 hours of infection. Following withdrawal of the oviposition plates, each of the above cages were provided with a fresh food plate (60 mm diameter) for 8 hours. From each of these plates (each coming from a single cage), three samples of 100 eggs each was picked using a moist paint brush and arranged onto the surface of three separate agar plates (90 mm Petri plates, 1.5% agar). These plates were incubated under standard laboratory maintenance conditions, and 48 hours later, the number of eggs on each agar plate that had hatched were visually counted using a light stereo microscope (Zeiss Stemi 2000) under 2.5X × 10X magnification. Hatchability was determined for each agar plate by dividing the number of eggs that hatched by the total number of eggs that were placed on the surface of that plate and was used as unit of replication. Altogether from each cage 3 × 100 eggs/treatment/population/block were scanned for hatchability.

### 2.6. Egg-to-adult development time and viability, and dry body weight

After 40 generation of forward selection, we tested for the effect of selection history on egg-to-adult development time and viability, and dry body weight at eclosion, of the flies from the EPN populations.

Fresh food plates with excess live yeast paste were provided to the standardised population cages (E_i_, P_i_, an N_i_) for 48 hours. Following this, a fresh food plate with live yeast paste (yeast paste placed at the centre of the plate and with space around the circumference for females to lay eggs) was provided to each cage for 6 hours. This plate was followed up with two more similarly yeasted food plates, each for a 1-hour window. This was done to encourage the females to lay any stored eggs. After this a fresh food plate was provided to each cage for females to lay eggs on for an hour, and eggs were collected using a light stereo microscope (Zeiss Stemi 2000) under 2.5X × 10X magnification and distributed into food vials (with 8 ml of banana-jaggery-yeast food) at an exact density of 70 eggs per vial. 10 vials were set up for each population (E_i_, P_i_, an N_i_), and blocks were handled on separate days. These vials were incubated under standard maintenance conditions, and when flies started eclosing, freshly eclosed flies were transferred to empty glass vials every 4 hours, and frozen at -20 °C for further processing; this was done till the very last pupae had eclosed. The storage vials were labelled so as to preserve parent vial, population, and block identities. The flies eclosed at each time window from each vial was later scored visually using a light stereo microscope (Zeiss Stemi 2000) under 2.5X × 10X magnification to enumerate the total number and sex of the flies eclosed in that time window. Therefore, for each vial, the data was available for the number of males and females that eclosed at each time window, and the total number of flies that eclosed out of the vial. The median development time was calculated as the time taken by half of flies of each sex to eclose (starting for the time of oviposition), and viability was calculated by dividing the total number of flies eclosed by the number of eggs seeded in the vial (which was 70 eggs). After enumeration, the flies were put back into -20 °C storage for further use.

Flies preserved from the development time assay were used for measuring dry body weight at eclosion. All flies eclosing out of a single parent vial were pooled together; hence there were 10 pools of flies per population per block. From each pool, 5 females and 5 males were randomly sampled and placed in 1.5 ml micro-centrifuge tubes (MCTs); females and males were placed in separate MCTs. Therefore, each vial used in the development time assay yielded one MCT with 5 males and one MCT with 5 females. These MCTs were dry heated in a hot air oven for 48 hours at 60 °C to eliminate all moisture. The flies were then weighed using Sartorius weighing balance (model CPA225D, least count 0.01mg). Individual blocks were handled on separate days. Dry body weight was measured for only 3 blocks because samples of block 2 was lost in handling.

### 2.7. Starvation resistance

After 37-38 generations of forward selection, we measured resistance to starvation of the EPN flies, and tested if selection history and infection status had an effect on starvation resistance.

Eggs were collected from standardised population cages at an approximate density of 70 eggs per vial. 20 such vials were collected for each population (E_i_, P_i_, an N_i_), and reared under standard maintenance conditions. Adults were housed in the rearing vials until 12^th^ day PEL. By this time all flies were sexually mature and have mated at least once inside the rearing vials itself. On 12^th^ day PEL, flies from each population (E_i_, P_i_, an N_i_) were randomly assigned to three treatments:

*infected* treatment: 50 females and 50 males were randomly sampled from each population, and were infected with *E. faecalis* at OD_600_ = 0.8 under light CO_2_ anaesthesia;
*sham-infected* treatment: 50 females and 50 males were randomly sampled from each population, and sham-infected with sterile needle dipped in sterile 10 mM MgSO_4_ solution; and,
*uninfected* treatment: 50 females and 50 males were just randomly sampled under light CO_2_ anaesthesia, and not subjected to any further manipulation.

After being subjected to treatments, the flies were housed in vials containing 2 ml 1.5% non-nutritive agar gel, at a density of 10 females (or, males) per vial. The sexes were housed separately. The presence of agar gel in the vials ensured that the flies had *ad libitum* access to water during the course of the starvation assay. Individual blocks were assayed upon on separate days. In total, 5 vials/sex/treatment/population/block were set up for this assay. The vials were monitored every 6-8 hours to record the number of dead flies, until the last fly perished. Surviving flies were transferred to fresh agar vials every 48 hours.

### 2.8. Desiccation resistance

After 38-39 generations of forward selection, we measured resistance to starvation of the EPN flies, and tested if selection history and infection status had an effect on starvation resistance.

A set-up identical to the starvation resistance assay was utilised to test for desiccation resistance; the only difference was that the flies, after being subjected to their respective treatments, were housed in empty vials (no food or agar gel). Additionally, 5 gm silica beads were placed in each vial (above the cotton plug; no direct contact between flies and silica), and the mouth of the vial was sealed off with parafilm tape, to eliminate moisture from the vials. Individual blocks were assayed upon on separate days. In total, 5 vials/sex/treatment/population/block were set up for this assay. The vials were monitored every 1.5 hours to record the number of dead flies, until the very last fly perished.

### 2.9. Larval competitive assay

After 40 generations of forward selection, we assayed larval competitive ability as a proxy of intra-specific competition. The larval competitive assay between focal (E_i_, P_i_, and N_i_) and competitor (PJB_w_) population was measured under two ratios. First, where focal and competitor were present at equal density (1 focal : 1 competitor), and second, where focal was present in one-third density to the competitor (1 focal : 3 competitor). The focal populations had wild-type, red eye colour and competitor had a homozygous-recessive white eye colour marker. The total density of eggs and available food volume in each vial was kept constant, that is, 100 eggs in 8ml of standard banana-jaggery-yeast food. For 1 focal : 1 competitor set-up, 50 eggs of focal population and 50 eggs of competitor population was placed in standard vials (25mm diameter × 90 mm height). Ten such vials were set for each focal population. Similarly, for 1 focal: 3 competitor set-up, 25 eggs of focal population were placed along with 75 eggs of competitor population in standard vial. Ten such replicate vials for each focal population were set up.

To set-up the larval competition assay, fresh food plates smeared with excess live yeast paste were provided to the standardised population cages (E_i_, P_i_, an N_i_) for 48 hours. Post 48 hours, a fresh food plate with live yeast paste (yeast paste placed at the centre of the plate and with space around the circumference for females to lay eggs) was provided to each cage for 6 hours. This plate was followed up with two more similarly yeasted food plates, each for a 1-hour window. This was done to encourage the females to lay any stored eggs. After this a fresh food plate was provided to each cage for females to lay eggs on for two hours, and eggs were collected at required densities using a light stereo microscope (Zeiss Stemi 2000) under 2.5X × 10X magnification and placed into food vials. 10 vials were set up for each population (E_i_, P_i_, an N_i_) and each competitive ratio (1focal: 1competitor, and 1focal: 3competitor). Blocks were handled on separate days. These bunches were reared under standard laboratory conditions until 12th day. On 13^th^ day, culture vials having adult flies were frozen in -20°C and later scored according to the eye colour. We scored adult survivors, as proxy for fitness of larvae to reach adulthood. Altogether 10 vials/population/ratio/block were assayed. Competitive index for the focal populations for each vial was calculated as,

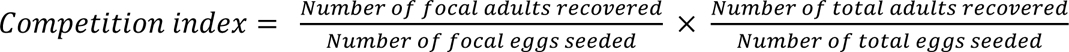

This competitive index for each vial was used as the unit of analysis.

### 2.1.0 Statistical analysis

All analysis was performed using R statistical software, version 4.1.0 (R Core Team 2021). Mixed-effect cox-proportional hazard models were fitted to the survival data (from post- infection survival, starvation resistance, and desiccation resistance assays) using the *coxme* function of the “coxme” package (Therneau 2020), and the confidence intervals for these models were calculated using *confint* function of the base R package. Survival curves were plotted using the *ggsurvplot* function of the “survminer” package (Kassambara et al 2021) after modelling the data using *survfit* function from the “survival” package (Therneau 2021).

For the analysis of data from the post-infection survival assay (first 96 hours following infection) we first modelled the total data as: survival ∼ infection treatment + (1|block), to test for the effect of infection treatment on survival, where ‘infection treatment’ was considered as a fixed factor and block as a random factor. Next, we modelled the data from infected flies only to test for the effect of selection history and sex on post-infection survival: survival ∼ selection + sex + selection:sex + (1|block), where ‘selection’ and ‘sex’ were considered as fixed factors and ‘block’ as a random factor.

Survival data from longevity assay (survival from 96 hours post-infection and onwards), starvation resistance assay, and desiccation resistance assay were analysed using the following model: survival ∼ selection + infection treatment + selection:infection treatment + (1|block), where ‘selection regime’ and ‘infection treatment’ were considered as fixed factors and ‘block’ as random factor. The data for each sex was analysed separately.

Data from life-history traits were modelled using mixed-effects general linear models (*lmer* function from “lmerTest” package; Kuznetsova et al 2017) and then subjected to type-III analysis of variance (ANOVA; *anova* function from base R package) for significance tests. Pairwise comparisons wherever necessary was done using Tukey’s HSD (*lsmeans* function from “emmeans” package; Lenth 2021). The mixed-effects general linear models used were: fecundity ∼ selection + infection treatment + selection:infection treatment + (1|block), hatchability ∼ selection + infection treatment + selection:infection treatment + (1|block), development time ∼ selection + sex + selection:sex + (1|block), viability ∼ selection + (1|block), body weight ∼ selection + sex + selection:sex + (1|block), and larval competition index ∼ selection + (1|block).

## 3. RESULTS

### 3.1. Response to selection, longevity, and fecundity

Survival during the pre-selection window (first 96 hours following infection) was affected by infection treatment. Both sham-infected (hazard ratio, 95% confidence interval: 11.744, 5.704- 24.179) and infected (HR, 95% CI: 171.679, 85.735-343.776) flies perished more following handling (infection/sham-infection) compared to uninfected flies. Among infected flies, E (HR, 95% CI: 0.608, 0.522-0.708) flies died less compared to N flies, while P (HR, 95% CI: 0.956, 0.831-1.098) and N flies had similar mortality. Sex of the hosts or interaction between host sex and selection history had no effect on survival of infected flies (table 1.b).

**Table 1.** Output of mixed-effects Cox proportional hazards model for analysis of post-infection survival (data from first 96 hours following infection). Hazard ratios are relative to the default level for each factor which is set to 1. The default level for “Treatment” is ’Uninfected’, the default level for “Selection” is ’N’, and the default level for “Sex” is ’Females’. Significant effects are marked in bold.

|  | Hazards Ratio | Lower CI (95%) | Upper CI (95%) | Z | p-value | Variance (for random factor) |
| --- | --- | --- | --- | --- | --- | --- |
| (a) Effect of infection treatments on overall survival |  |  |  |  |  |  |
| Treatment *Sham-infected* | **11.7441** | **5.704236** | **24.17919** | **6.69** | **2.3e-11** |  |
| Treatment *Infected* | **171.6795** | **85.735467** | **343.77657** | **14.52** | **0.0e+00** |  |
| Block |  |  |  |  |  | 0.1007759 |
| (b) Effect of selection history and sex on post-infection survival |  |  |  |  |  |  |
| Selection *P* | 0.9556693 | 0.8313593 | 1.0985669 | -0.64 | 5.2e-01 |  |
| **Selection *E*** | **0.6078106** | **0.5218545** | **0.7079248** | **-6.40** | **1.6e-10** |  |
| Sex *Males* | 0.9143132 | 0.7944967 | 1.0521990 | -1.25 | 2.1e-01 |  |
| Selection *P* :<br>Sex *Males* | 1.0065507 | 0.8239498 | 1.2296190 | 0.06 | 9.5e-01 |  |
| Selection *E* :<br>Sex *Males* | 0.8391703 | 0.6703517 | 1.0505036 | -1.53 | 1.3e-01 |  |
| Block |  |  |  |  |  | 0.1141784 |

Longevity of flies from the selection window onwards (day 5 following infection and onwards) was significantly affected by infection treatment in case of both female and male flies. Both sham-infected (HR, 95% CI: 1.437, 1.237-1.670) and infected (HR, 95% CI: 7.790, 6.654- 9.120) females died faster compared to uninfected females. Similarly, both sham-infected (HR, 95% CI: 1.219, 1.050-1.415) and infected (HR, 95% CI: 4.767, 4.109-5.530) males died significantly faster compared to uninfected males. In case of both females (table 2.a) and males (table 2.b), selection history had no effect on longevity of flies.

**Table 2.** Output of mixed-effects Cox proportional hazards model for analysis of longevity (data from day 5 post-infection onwards). Hazard ratios are relative to the default level for each factor which is set to 1. The default level for “Treatment” is ’Uninfected’, and the default level for “Selection” is ’N’. Significant effects are marked in bold.

| Factors | Hazards Ratio | Lower CI (95%) | Upper CI (95%) | Z | p-value | Variance (for random factor) |
| --- | --- | --- | --- | --- | --- | --- |
| (a) Females |  |  |  |  |  |  |
| Selection *P* | 1.1551639 | 0.9969687 | 1.3384609 | 1.92 | 5.5e-02 |  |
| Selection *E* | 1.0767328 | 0.9308278 | 1.2455080 | 1.00 | 3.2e-01 |  |
| Treatment *Sham* | **1.4373292** | **1.2368143** | **1.6703519** | **4.73** | **2.2e-06** |  |
| Treatment *Infected* | **7.7901587** | **6.6542407** | **9.1199847** | **25.53** | **0.0e+00** |  |
| Selection *P* : Treatment *Sham* | **0.6824614** | **0.5519484** | **0.8438355** | **-3.53** | **4.2e-04** |  |
| Selection *E* : Treatment *Sham* | 0.9452391 | 0.7664630 | 1.1657144 | -0.53 | 6.0e-01 |  |
| Selection *P* : Treatment *Infected* | 0.9865656 | 0.8009505 | 1.2151957 | -0.13 | 9.0e-01 |  |
| Selection *E* : Treatment *Infected* | 0.9557911 | 0.7814021 | 1.1690993 | -0.44 | 6.6e-01 |  |
| Block (Random) |  |  |  |  |  | 0.01164269 |
| (b) Males |  |  |  |  |  |  |
| Selection *P* | 1.0021483 | 0.8663334 | 1.159255 | 0.03 | 0.9800 |  |
| Selection *E* | 1.0374557 | 0.8957662 | 1.201557 | 0.49 | 0.6200 |  |
| Treatment *Sham* | **1.2189010** | **1.0496558** | **1.415435** | **2.60** | **0.0094** |  |
| Treatment *Infected* | **4.7670378** | **4.1092087** | **5.530176** | **20.61** | **0.0000** |  |
| Selection *P* : Treatment *Sham* | 1.0102395 | 0.8205032 | 1.243851 | 0.10 | 0.9200 |  |
| Selection *E* : Treatment *Sham* | 1.1893644 | 0.9644225 | 1.466772 | 1.62 | 0.1000 |  |
| Selection *P* : Treatment *Infected* | 0.9119939 | 0.7448162 | 1.116695 | -0.89 | 0.3700 |  |
| Selection $E$ :<br>Treatment<br>*Infected* | 1.0874873 | 0.8922925 | 1.325382 | 0.83 | 0.4100 | |
| Block<br>(Random) |  |  |  |  |  | 0.03896682 |

Fecundity (per-female) during the selection window (between 96^th^ and 114^th^ hour following infection) was not affected by either selection history or infection treatment of the flies (table 3.a). Hatchability of the eggs laid during this same period was also not affected by either selection history or infection treatment of the flies (table 3.b).

**Table 3.** Type III analysis of variance (ANOVA) output for adult life-history traits. Significant effects are marked in bold.

| Factors | SS | MS | df | Residual df | F value | p value |
| --- | --- | --- | --- | --- | --- | --- |
| (a) Female fecundity |  |  |  |  |  |  |
| Selection | 6.2074 | 3.1037 | 2 | 36 | 1.2708 | 0.2929 |
| Treatment | 7.0083 | 3.5041 | 2 | 36 | 1.4347 | 0.2515 |
| Selection × Treatment | 2.1134 | 0.5284 | 4 | 36 | 0.2163 | 0.9276 |
| (b) Hatchability |  |  |  |  |  |  |
| Selection | 0.00111321 | 0.00055661 | 2 | 104 | 1.5011 | 0.2277 |
| Treatment | 0.00017956 | 0.00008978 | 2 | 104 | 0.2421 | 0.7854 |
| Selection × Treatment | 0.00089324 | 0.00022331 | 4 | 104 | 0.6022 | 0.6619 |

### 3.2. Development time, egg-to-adult viability, and body weight at eclosion

Sex had a significant effect on egg-to-adult development time (F_1,232_: 1656.81, p = 1.219 e-05), with females eclosing earlier than males. Selection history or selection history × sex interaction had no effect on development time (table 4.a). Selection history also had no effect on egg-to-adult viability (table 4.b).

**Table 4.**
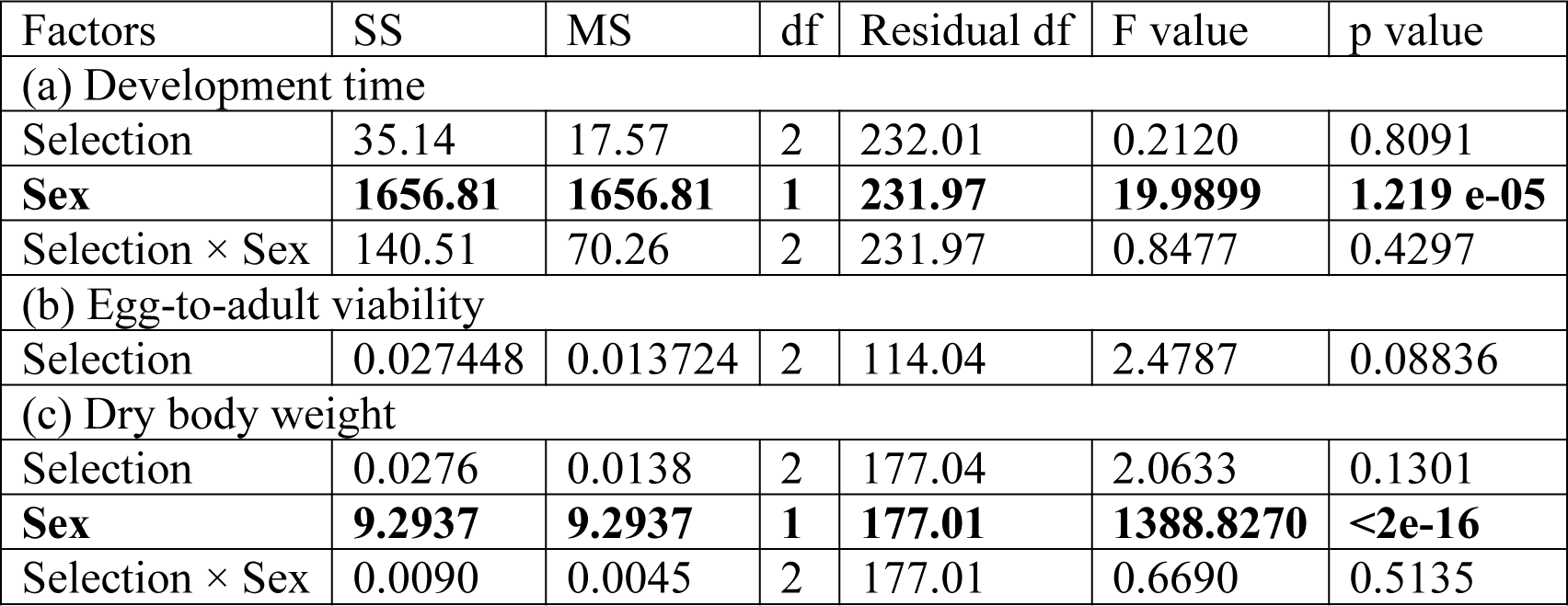
Type III analysis of variance (ANOVA) output for juvenile life-history traits. Significant effects are marked in bold.

Dry body weight at eclosion was significantly affected by sex (F_1,177_: 9.294, p < 2 e-16), with females having greater weight than males. Selection history or selection history × sex interaction had no effect on development time (table 4.c).

### 3.3. Starvation and desiccation resistance

Survival of female flies subjected to starvation was affected by infection treatment: infected females (HR, 95% CI: 2.418, 1.975-2.959) died faster compared to uninfected females, while sham-infected (HR, 95% CI: 1.080, 0.881-1.325) females and uninfected females died at a similar rate. Starvation resistance of female flies was not affected by selection history (table 5.a). Survival of male flies subjected to starvation was similarly affected by infection treatment: infected males (HR, 95% CI: 2.429, 1.995-2.958) died faster compared to uninfected males, while sham-infected (HR, 95% CI: 1.021, 0.837-1.246) males and uninfected males died at a similar rate. Starvation resistance of male flies was additionally affected by selection history: pooling all infection treatments together, E males (HR, 95%CI: 0.814, 0.666-0.993) perished due to starvation significantly later than N males; there was no significant difference between P and N males (table 5.b).

**Table 5.** Output of mixed-effects Cox proportional hazards model for analysis of starvation resistance data (survival under starved conditions). Hazard ratios are relative to the default level for each factor which is set to 1. The default level for “Treatment” is ’Uninfected’, and the default level for “Selection” is ’N’. Significant effects are marked in bold.

| Factors | Hazards Ratio | Lower CI (95%) | Upper CI (95%) | Z | p-value | Variance (for random factor) |
| --- | --- | --- | --- | --- | --- | --- |
| (a) Females |  |  |  |  |  |  |
| Selection *P* | 1.1751116 | 0.9609941 | 1.4369362 | 1.57 | 0.12000 |  |
| Selection *E* | 1.1577567 | 0.9451722 | 1.4181550 | 1.42 | 0.16000 |  |
| Treatment *Sham* | 1.0804146 | 0.8811486 | 1.3247433 | 0.74 | 0.46000 |  |
| **Treatment Infected** | **2.4178591** | **1.9753521** | **2.9594940** | **8.56** | **0.00000** |  |
| Selection *P* : Treatment *Sham* | 0.8232719 | 0.6196202 | 1.0938582 | - 1.34 | 0.18000 |  |
| Selection *E* : Treatment *Sham* | 1.1580500 | 0.8705147 | 1.5405595 | 1.01 | 0.31000 |  |
| **Selection *P* : Treatment Infected** | **0.6032435** | **0.4551948** | **0.7994440** | **- 3.52** | **0.00043** |  |
| **Selection *E* : Treatment Infected** | **0.7322360** | **0.5524562** | **0.9705195** | **- 2.17** | **0.03000** |  |
| Block (Random) |  |  |  |  |  | 0.1433253 |
| (b) Males |  |  |  |  |  |  |
| Selection *P* | 0.8863760 | 0.7276611 | 1.0797092 | - 1.20 | 0.230 |  |
| **Selection *E*** | **0.8135374** | **0.6664097** | **0.9931474** | **- 2.03** | **0.043** |  |
| Treatment *Sham* | 1.0212526 | 0.8369739 | 1.2461043 | 0.21 | 0.840 |  |
| **Treatment Infected** | **2.4291004** | **1.9948018** | **2.9579523** | **8.83** | **0.000** |  |
| Selection *P* : Treatment *Sham* | 1.2696626 | 0.9583504 | 1.6821018 | 1.66 | 0.096 |  |
| Selection *E* : Treatment *Sham* | 1.0708688 | 0.8071027 | 1.4208353 | 0.47 | 0.640 |  |
| Selection *P* : Treatment *Infected* | 1.0807332 | 0.8175647 | 1.4286139 | 0.55 | 0.590 |  |
| Selection $E$ :<br>Treatment<br>*Infected* | 0.9888991 | 0.7474084 | 1.3084165 | -<br>0.08 | 0.940 | |
| Block<br>(Random) |  |  |  |  |  | 0.02744811 |

Survival of female flies subjected to desiccation was affected by infection treatment, with both sham-infected (HR, 95% CI: 2.242, 1.838-2.734) and infected (HR, 95% CI: 2.117, 1.736- 2.583) females dying earlier than uninfected females. Selection history also had a significant effect on female desiccation resistance, with P females (HR, 95%CI: 1.242, 1.020-1.512) dying earlier than N females; there was no difference between mortality rate of E and N females (table 6.a). Survival of male flies subjected to desiccation was similarly affected by infection treatment, with both sham-infected (HR, 95% CI: 2.138, 1.754-2.606) and infected (HR, 95% CI: 2.463, 2.013-3.014) males dying earlier than uninfected males. Selection history also had a significant effect on male desiccation resistance, with E males (HR, 95%CI: 0.779, 0.640- 0.948) dying later than N females; there was no difference between mortality rate of P and N males (table 6.a).

**Table 6.**
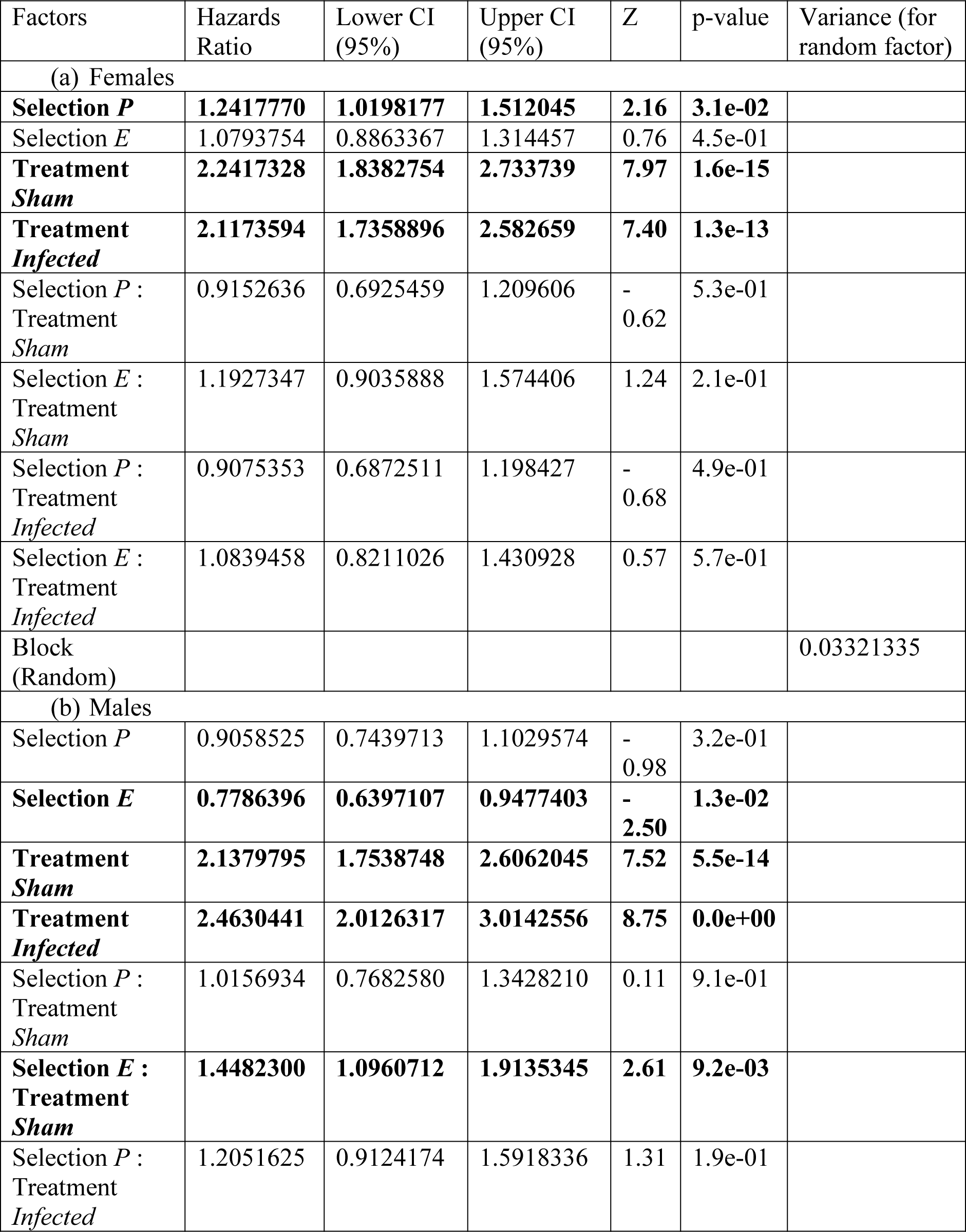
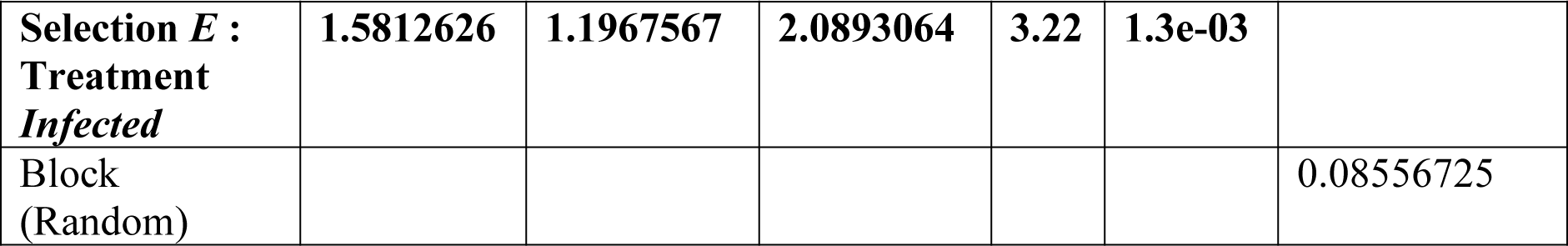
Output of mixed-effects Cox proportional hazards model for analysis of desiccation resistance data (survival under desiccated conditions). Hazard ratios are relative to the default level for each factor which is set to 1. The default level for “Treatment” is ’Uninfected’, and the default level for “Selection” is ’N’. Significant effects are marked in bold.

### 3.4. Intra-specific (larval) competition

Selection history had a significant effect on competitive index (see Methods for calculation of competitive index) irrespective of the infection intensity (ratio of focal and competitor eggs at the beginning of the assay; table 7). At low competition intensity (1 focal:1 competitor), P populations had a significantly higher competitive index than both N (t-ratio: -3.769, df: 109, p = 0.0008) and E (t-ratio: 3.685, df: 109, p = 0.0010) populations (post-hoc pairwise comparison using Tukey’s HSD). At high competition intensity (1 focal:3 competitor), E populations exhibited a significantly higher competitive index than both N (t-ratio: -2.405, df: 104, p = 0.0468) and P (t-ratio: -2.591, df: 104, p = 0.0292) populations (post-hoc pairwise comparison using Tukey’s HSD).

**Table 7.** Type III analysis of variance (ANOVA) output for larval competitive ability. Significant effects are marked in bold.

| Factors | SS | MS | df | Residual df | F value | p value |
| --- | --- | --- | --- | --- | --- | --- |
| (a) 1 focal : 1 competitor |  |  |  |  |  |  |
| **Selection** | **0.51269** | **0.25634** | **2** | **108** | **9.7583** | **0.0001271** |
| (b) 1 focal : 3 competitor |  |  |  |  |  |  |
| **Selection** | **0.36332** | **0.18166** | **2** | **102.06** | **4.2513** | **0.01685** |

## 4. DISCUSSION

### 4.1. Response to selection

We selected *Drosophila melanogaster* populations for increased post-infection survival when adults are infected with *Enterococcus faecalis*. After 35 generations of forward selection, flies of the selected populations (E populations) exhibited a marked reduction in post-infection mortality compared to flies from the control populations (P and N populations), indicating a successful response to selection (figure 1). Susceptibility to infection by *E. faecalis* was not determined by sex of the host in either the control or the selected populations (figure 1).

**Figure 1.**
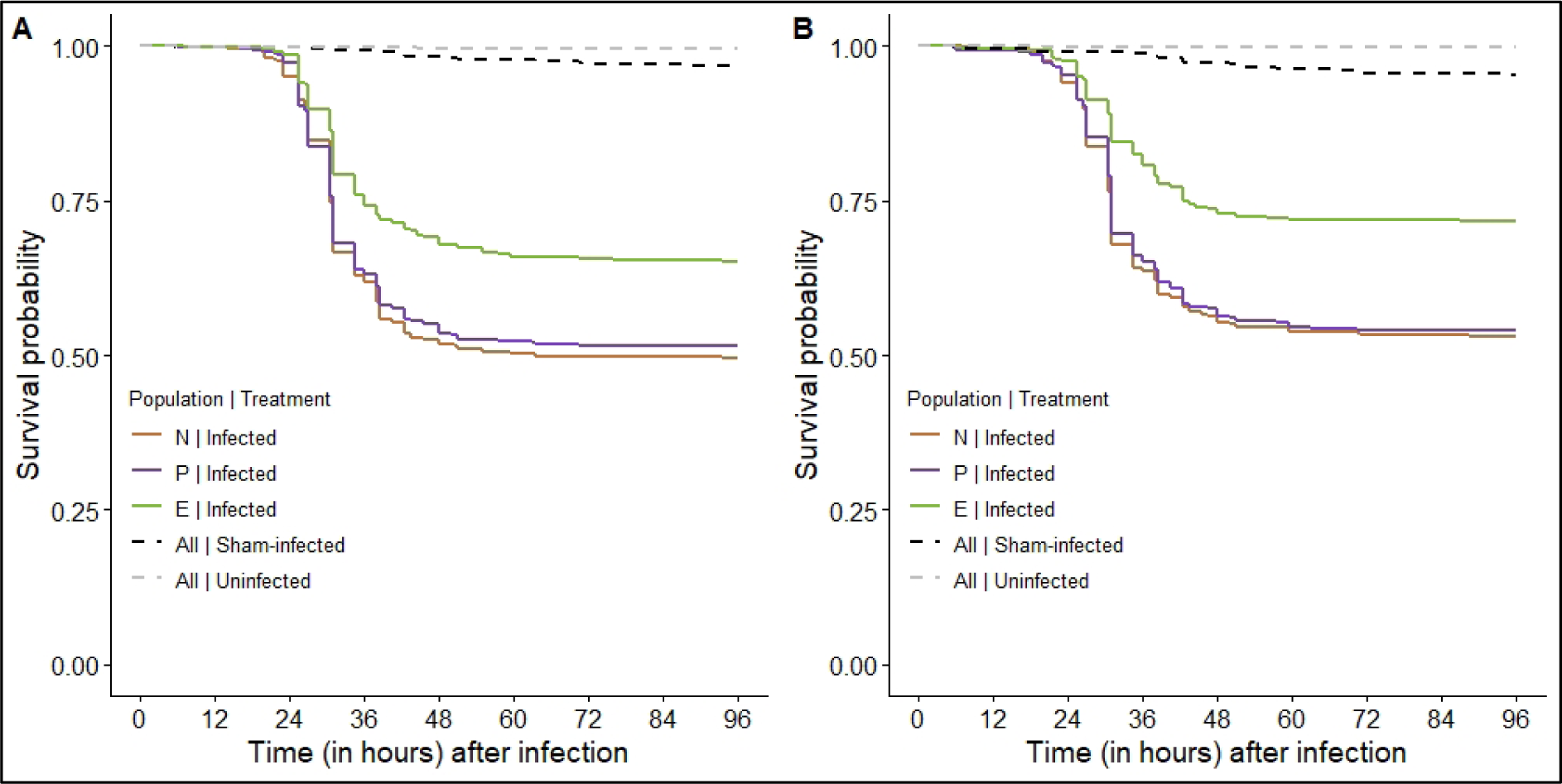
Post-infection survival of flies from EPN populations for the first 96 hours following infection: (A) females, and (B) males.

**Figure 2.**
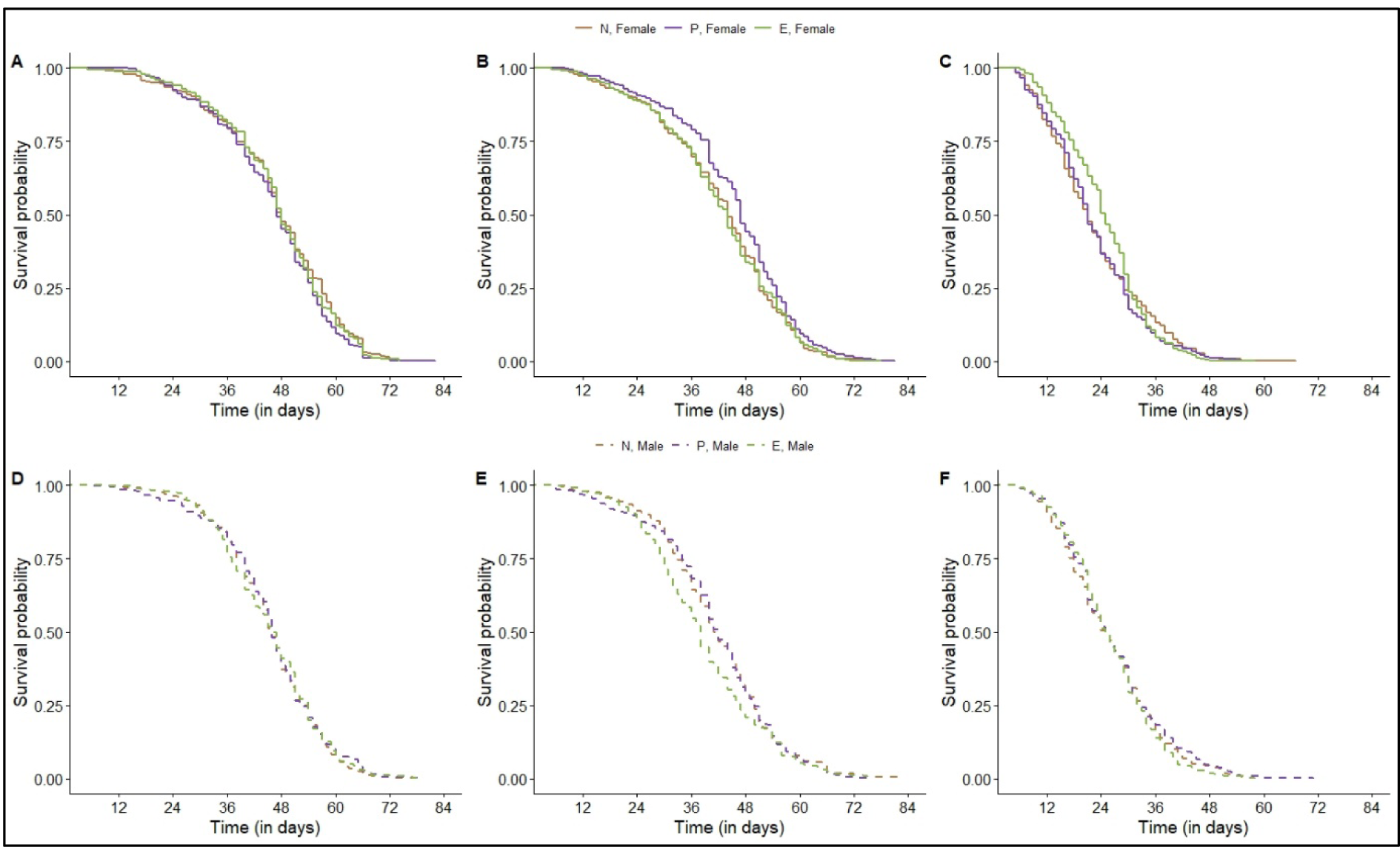
Life-time post-infection survival (longevity) of flies from EPN populations from day 5 post-infection onwards: (A) uninfected females, (B) sham-infected females, (C) infected females, (D) uninfected males, (E) sham-infected males, and (F) infected males.

### 4.2. Juvenile life-history traits

Egg-to-adult development time and viability (along with adult dry body weight at eclosion) was measured after 40 generations of forward selection. Egg-to-adult development time in the EPN populations was not affected by selection history (figure 4.A). Our results are similar to those obtained by Faria et al (2015) and Gupta et al (2016), both of whom selected flies to survive better following infection with *Pseudomonas entomophila* and found no effect of selection history on development time. Flies evolved to better survive infection with *Pseudomonas aeruginosa* have a shorter development time compared to their controls (Ye et al 2009). Egg-to-adult viability in the EPN populations was also not affected by selection history (figure 4.B), similar to reports from flies selected using *P. entomophila* (Faria et al 2015, Gupta et al 2016). Flies selected using *P. aeruginosa* exhibit reduced egg-to-adult viability (Ye et al 2009); it must be noted that Ye et al (2009) referred to their assay as *egg viability*, but their protocol suggests that what was measured was indeed egg-to-adult viability, and not the viability of eggs only. Put together these results suggest that the effect of evolving increased defense against bacterial pathogens (in the adult stage) on juvenile life history traits is determined by the identity of the pathogen used for selection. In the EPN populations, females developed faster than males (figure 4.A), which is common in *D. melanogaster* studies (reviewed in Prasad and Joshi 2003).

### 4.3. Adult life-history traits

Dry body weight at eclosion for adults was influenced by sex of the flies, with females being heavier than males, but within each sex there was no observable effect of selection history (figure 4.C). All three previous experimental evolution studies using bacterial pathogens have reported similar results (Ye et al 2009, Faria et al 2015, Gupta et al 2016).

Longevity and fecundity, with and without infection, was measured after 35 generations of forward selection. Selection history did not have any effect on female fecundity (figure 3.A) and hatchability of the eggs laid (figure 3.B) in the EPN populations. We define hatchability as the proportion of eggs that produced a living larva within forty hours of being laid. All three previous experimental evolution studies using bacterial pathogens have reported similar results (Ye et al 2009, Faria et al 2015, Gupta et al 2016). This suggests that evolution of increased defense against bacterial pathogens does not come at a cost of female reproductive capacity in *D. melanogaster* hosts. Interestingly, female fecundity (and hatchability of the eggs laid) was also unaffected by host infection status: females of all populations had comparable fecundity irrespective whether they were subjected to infection, sham-infection, or left uninfected (figures 3.A and 3.B). Although this goes against the theoretical expectations (Lochmiller and Deerenberg 2000, Schmid-Hempel 2003, McKean et al. 2008), our results are in line with some of the previous studies that have demonstrated an apparent lack of change in fecundity when females are infected with bacteria *P. entomophila*, *Lactococcus lactis*, and *Escherichia coli* (Kutzer and Armitage 2016, Kutzer et al 2018).

**Figure 3.**
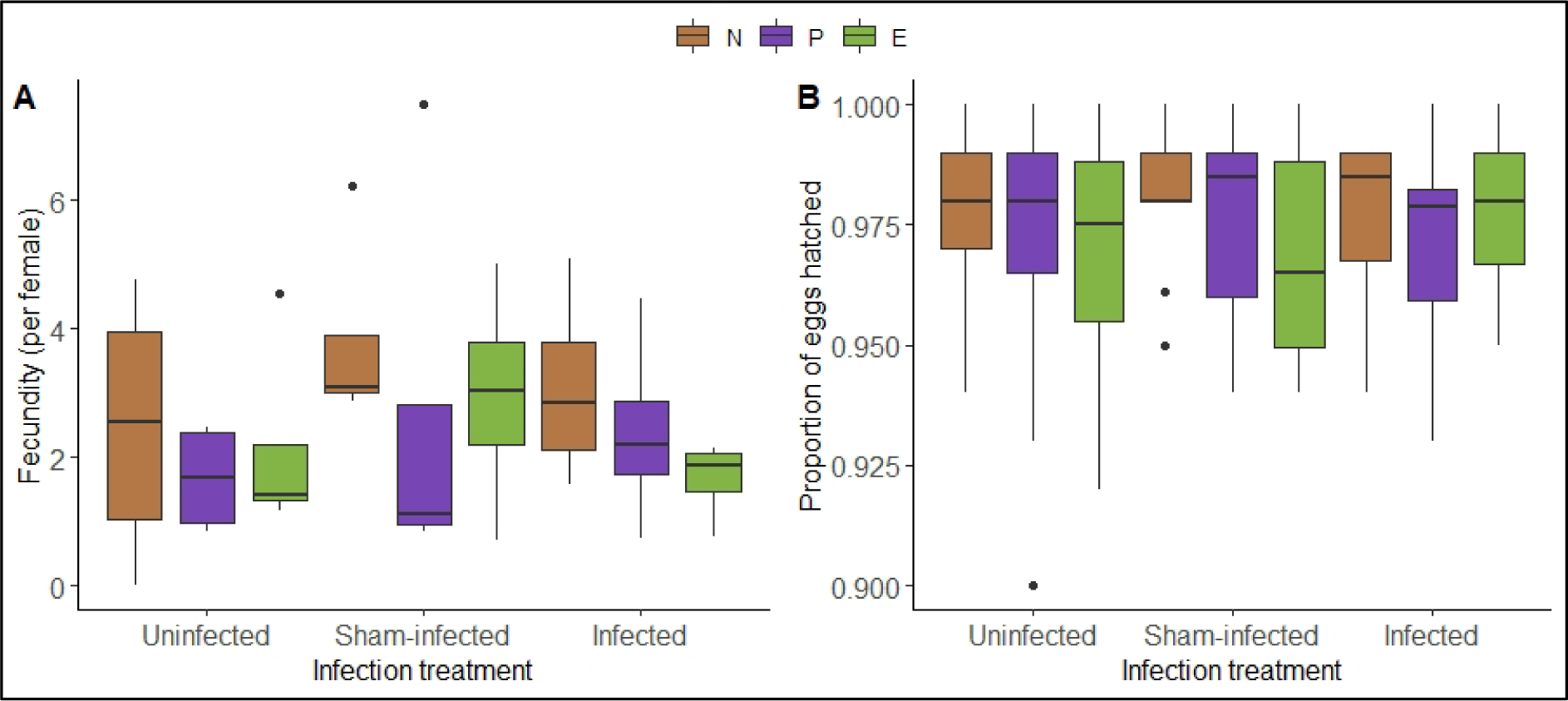
Reproductive output of females from EPN populations: (A) fecundity, and (B) egg hatchability.

**Figure 4.**
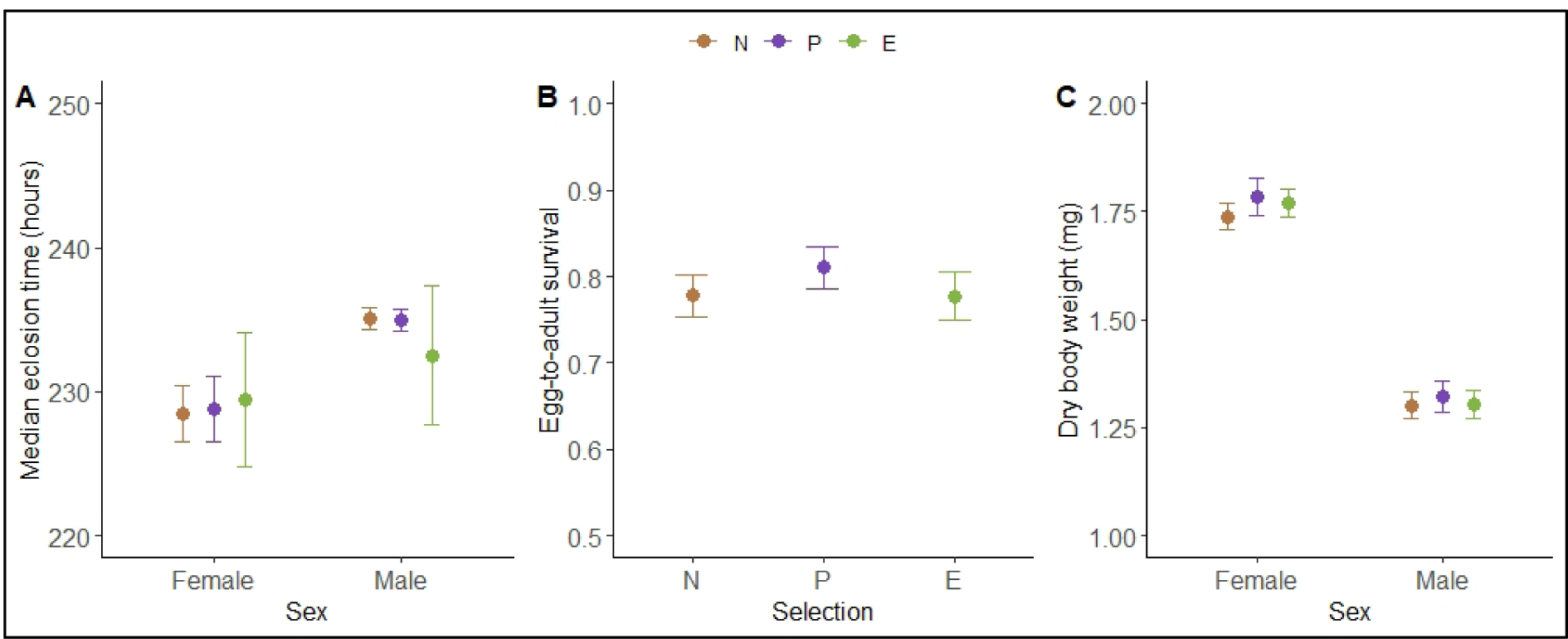
Juvenile life-history traits of flies from EPN populations: (A) egg-to-adult development time, (B) egg-to-adult survival, and (C) dry body weight at eclosion.

Selection history had no overall effect on longevity of either females or males (figure 2). Previous studies have shown that flies evolved to defend against *P. aeruginosa* have shorter life-span compared to control flies (Ye et al 2009), but that is not the case for flies evolved to defend against *P. entomophila* (Gupta et al 2016), suggesting that the consequences of evolving increased immunity on host life-span is pathogen specific. In case of both sexes, adult life-span was determined by the infection status of the flies: infected flies of both sexes survived less compared to their sham-/uninfected counterparts. Please note that in this context the infected flies represent such individuals who have survived the acute phase of infection. A few possible hypotheses, individually or together, can explain why survivors of acute infection die early compared to control flies. One possibility is that surviving the acute phase of infection implies mounting a successful immune defense, and the early death (reduced lifespan) is an associated cost, probably because of exhaustion of resources or permanent damage to the soma caused by the pathogen. Additionally, flies that have survived acute infection continue to harbor very low dose of pathogens in their system (chronic infection), and it takes a continuous and costly investment towards immune function to ensure that the pathogen load does not re-increase (Chambers et al. 2019). A third possibility is that survivors of acute infection die early because of the damage to their organs (immunopathology) caused by their own immune response (Khan et al 2017).

### 4.4. Response to abiotic stressors

Resistance to abiotic stressors was measured between 35-38 generations of forward selection. Similar to what was reported by Faria et al (2015) and Gupta et al (2016), we do not observe an increase in susceptibility to either starvation (figure 5) or desiccation (figure 6) in the selected populations (E populations) compared to the control populations (P and N populations). In fact, E population males are more resistant to both starvation and desiccation (when all infection treatments are pooled together) compared to males from N populations.

**Figure 5.**
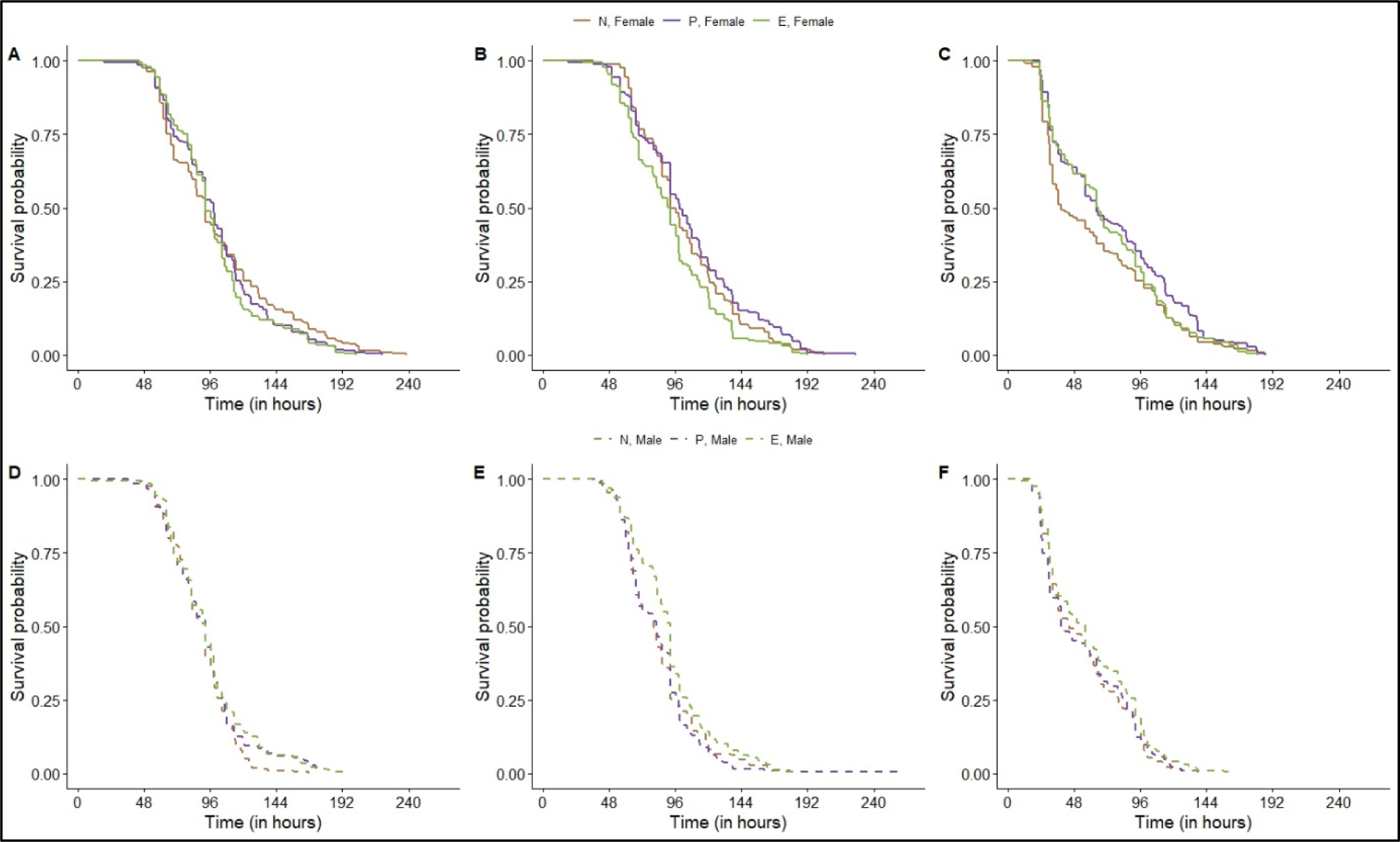
Starvation resistance (survival under starved conditions) of adult flies from EPN populations: (A) uninfected females, (B) sham-infected females, (C) infected females, (D) uninfected males, (E) sham-infected males, and (F) infected males.

**Figure 6.**
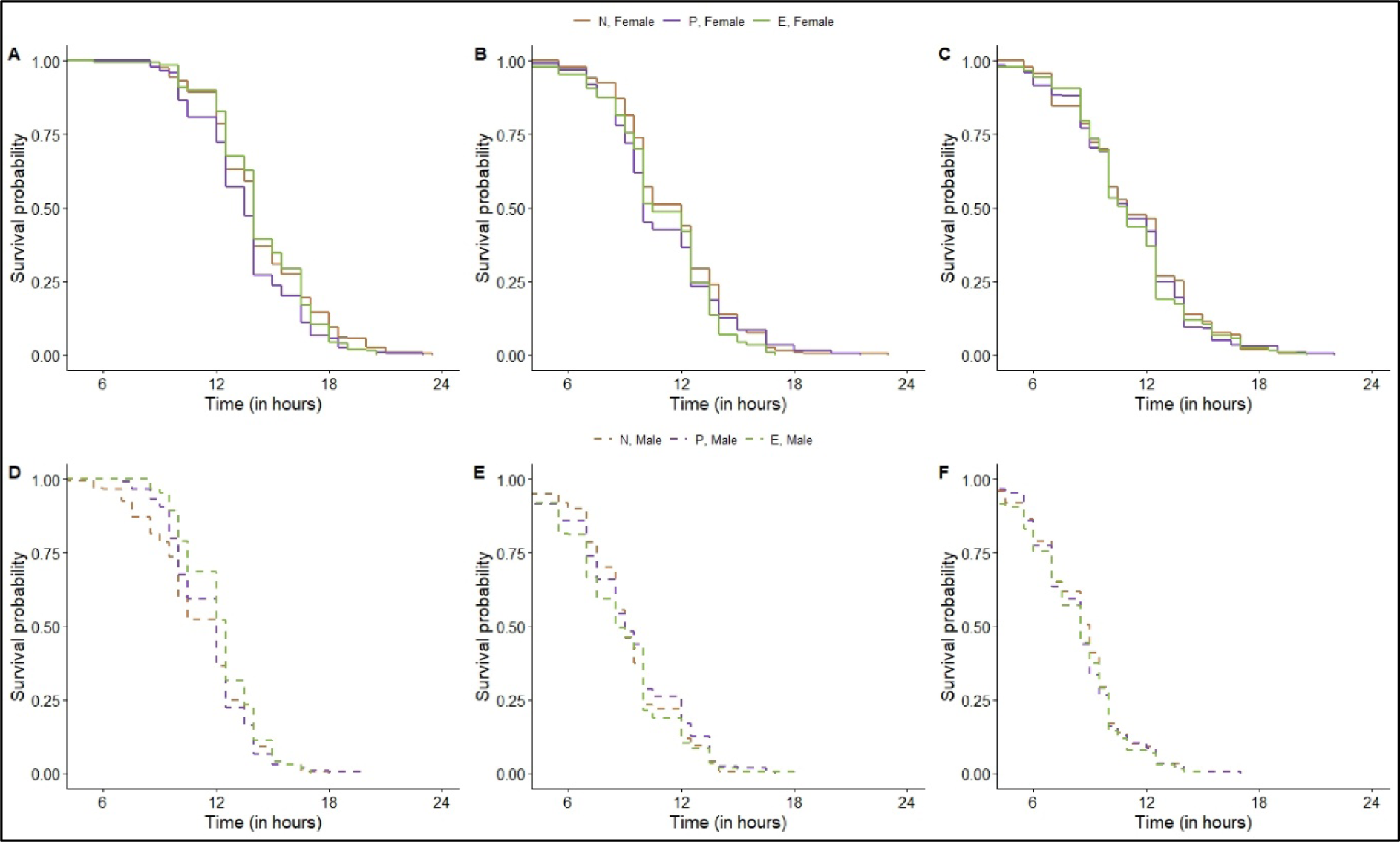
Desiccation resistance (survival under desiccated conditions) of adult flies from EPN populations: (A) uninfected females, (B) sham-infected females, (C) infected females, (D) uninfected males, (E) sham-infected males, and (F) infected males.

Infection status of the host had a significant effect on susceptibility to both biotic stressors. Infected flies, from all three selection regimes and both sexes, succumbed to starvation earlier compared to both sham-infected and uninfected flies, both of which perished at a comparable rate (figure 5). Early mortality of infected flies when starved might also be a manifestation of costs of mounting an immune response, similar to the case of adult lifespan. Previous authors have suggested a correlation between resistance to starvation and adult longevity in *Drosophila melanogaster* (reviewed in Prasad and Joshi 2003, Rion and Kawecki 2007), so similarity in observations from our longevity and starvation resistance assays is not surprising.

The effect of infection status on desiccation resistance were very different. Both infected and sham-infected flies (irrespective of sex and selection history) succumbed to desiccation before the uninfected flies; there was no discernable difference between the mortality rate of the infected and sham-infected flies (figure 6). One possible explanation for this observation is that the procedure for both infection and sham-infection involve pricking the flies with a fine needle, leading to a breach of the cuticle, which can lead to loss of haemolymph, and moisture in general. Rate of losing moisture is a major determinant of desiccation resistance in *Drosophila melanogaster* (reviewed in Prasad and Joshi 2003). Also, desiccation is a much faster acting stress compared to starvation, working at a time scale shorter than even the time taken by the pathogen to kill the flies. Flies start dying from infection around 18-20 hours post-infection, while even the most long-lived fly under desiccation stress doesn’t live till 24 hours. This might explain the lack of difference between mortality rate between the infected and sham-infected flies when subjected to desiccation.

### 4.5. Response to biotic stressor

We measured larval competitive ability, as a proxy of intra-specific competition, after 40 generations of forward selection. Larval competitive ability across both competition environment was affected by selection history (figure 7), but the difference between selected and control populations was not consistent across different competition environments. When the competition assay was run starting with equal numbers of eggs from focal and competitor populations, the E populations had a competitive index comparable to N populations, while a lower competitive index compared to P populations. When the assay was run starting with focal and competitor eggs in 1:3 ratio, E populations had a higher competitive index compared to both N and P populations. As of yet, we do not have an explanation for this discrepancy. Previous studies have shown that flies selected for resistance against larval parasitoids have reduced larval competitive ability, especially when resources are scarce (Kraaijeveld and Godfray 1997, Fellowes et al 1998). Our results might differ from theirs because of two notable differences: one, the pathogen/parasite used for selection are different (bacteria vs. parasitoid), and two, the life stage at which selection is applied are different (adults vs. larva).

**Figure 7.**
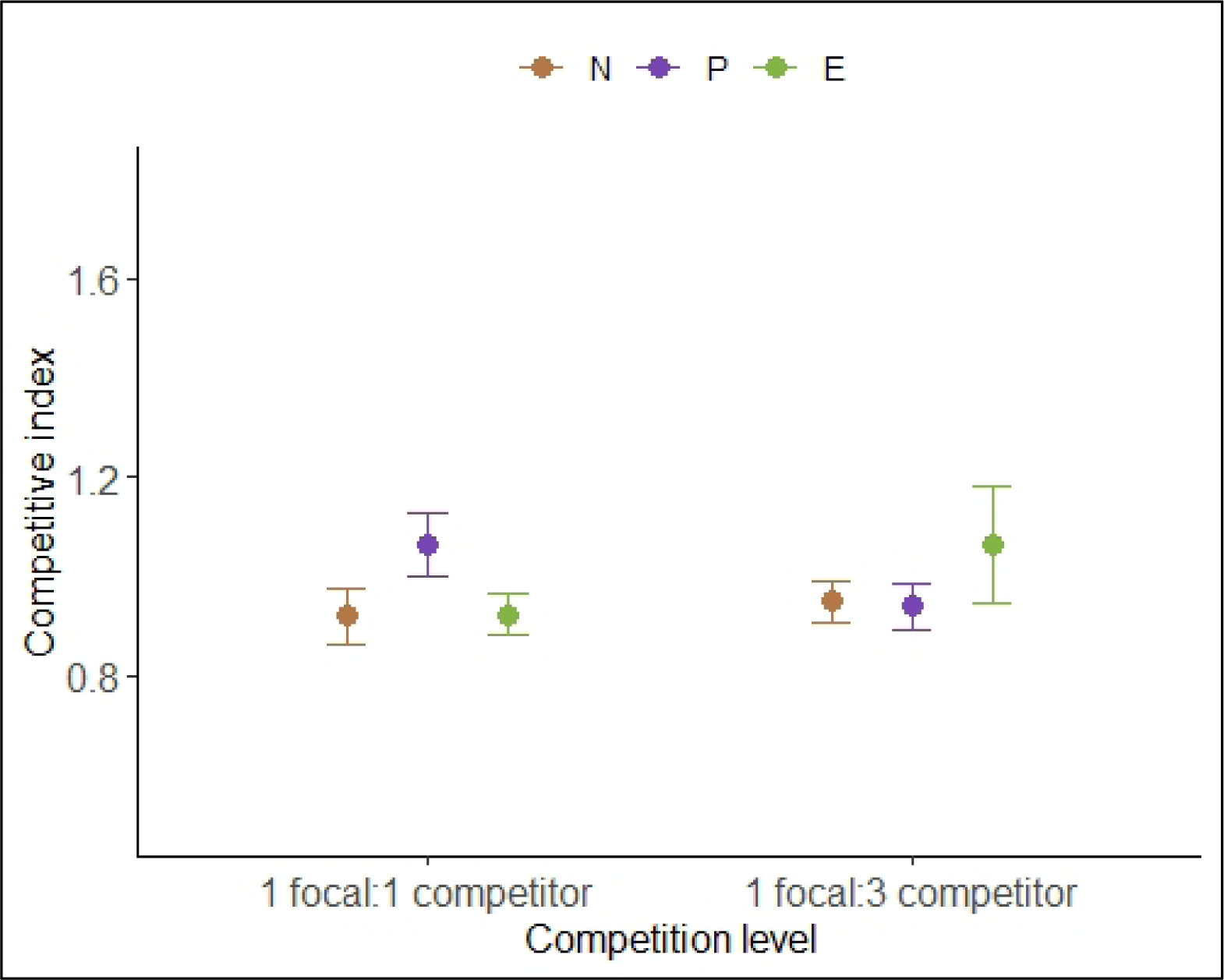
Larval competitive ability of EPN populations.

## 5. CONCLUSION

We tested for different types of costs of immunity: evolutionary costs of evolving an increased defense and physiological costs of mounting an immune defense, using replicate populations of *Drosophila melanogaster* selected for increased post-infection survival following infection with a Gram-positive bacterium, *Enterococcus faecalis*. We found no evidence of evolutionary costs: the selected population and control populations did not differ from one another in terms of trait values of life-history traits, either in the juvenile or in the adult stage. Selected populations also did not exhibit an increased susceptibility to abiotic stress. Put together with previous studies that have experimentally evolved fly populations for increased immunity against bacterial population (Ye et al 2009, Faria et al 2015, Gupta et al 2016, Ahlawat et al 2022), we propose that whether evolving increased defense comes at the cost of other organismal function depend on the bacterial pathogen used for selection. The cost of mounting an immune defense was specific to the trait under focus but did not differ across different selection histories. Infected flies exhibited shorter lifespan compared to uninfected flies, but there was no effect of infection status on female reproductive output. Resistance to starvation was also compromised in infected flies compared to uninfected flies. This suggests that physiological trade-offs between immune function and other organismal functions is not a universal expectation.

## Author contributions

Conceptualization and Methodology: AS, AKB, TH, NGP; Execution of experiments and data collection: AS, AKB, TH, BS, NB, AC; Statistical analysis: AS, AKB; Manuscript preparation: AS, AKB. Fund acquisition and supervision: NGP.

## Funding

This study was funded by IISER Mohali intramural funds and a research grant (no. BT/PR14278/BRB/10/1417/2015) from Dept. of Biotechnology, Govt. of India. AS was supported by Junior and Senior Research Fellowships from University Grants Commission, Govt. of India. AKB was supported by Junior and Senior Research Fellowships from Council of Scientific and Industrial Research, Govt. of India. TH and NB were supported by KVPY Fellowship, and BS was supported by INSPIRE Scholarship for Higher Education, both from Dept. of Science and Technology, Govt. of India.

## Acknowledgement

We thank Prof. B Lazzaro (Cornell University, Ithaca, USA) for providing us *Enterococcus faecalis*. We acknowledge Chinmay Krishna and Ruben Aju George for help during experiments.

## Conflict of interests

Authors have no competing interests to declare.

## Notes

### Competing Interest Statement

The authors have declared no competing interest.

## REFERENCES

1. Ahlawat, N., Maggu, K., Jigisha, Arun, M.G., Meena, A., Agarwala, A., Prasad, N.G., 2022. No major cost of evolved survivorship in *Drosophila melanogaster* populations coevolving with *Pseudomonas entomophila*. Proceedings of Royal Society B 289, 20220532. https://doi.org/10.1098/rspb.2022.0532

2. Brandt, S.M., Schneider, D.S., 2007. Bacterial infection of fly ovaries reduces egg production and induces local hemocyte activation. Developmental & Comparative Immunology 31, 1121–1130. https://doi.org/10.1016/j.dci.2007.02.003

3. Chambers, M.C., Jacobson, E., Khalil, S., Lazzaro, B.P., 2019. Consequences of chronic bacterial infection in *Drosophila melanogaster*. PLOS ONE 14, e0224440. https://doi.org/10.1371/journal.pone.0224440

4. Faria, V.G., Martins, N.E., Paulo, T., Teixeira, L., Sucena, É., Magalhães, S., 2015. Evolution of Drosophila resistance against different pathogens and infection routes entails no detectable maintenance costs. Evolution 69, 2799–2809. https://doi.org/10.1111/evo.12782

5. Fellowes, M.D.E., Kraaijeveld, A.R., Godfray, H.C.J., 1998. Trade–off associated with selection for increased ability to resist parasitoid attack in *Drosophila melanogaster*. Proceedings of the Royal Society of London. Series B: Biological Sciences 265, 1553– 1558. https://doi.org/10.1098/rspb.1998.0471

6. Gupta, V., Stewart, C.O., Rund, S.S.C., Monteith, K., Vale, P.F., 2017. Costs and benefits of sublethal Drosophila C virus infection. Journal of Evolutionary Biology 30, 1325–1335. https://doi.org/10.1111/jeb.13096

7. Gupta, V., Venkatesan, S., Chatterjee, M., Syed, Z.A., Nivsarkar, V., Prasad, N.G., 2016. No apparent cost of evolved immune response in *Drosophila melanogaster*. Evolution 70, 934–943. https://doi.org/10.1111/evo.12896

8. Hudson, A.L., Moatt, J.P., Vale, P.F., 2020. Terminal investment strategies following infection are dependent on diet. Journal of Evolutionary Biology 33, 309–317. https://doi.org/10.1111/jeb.13566

9. Kassambara, A., Kosinski, M., and Biecek, P., (2021). survminer: 676 Drawing Survival Curves using ’ggplot2’. R package version 0.4.9.

10. Khan, I., Agashe, D., Rolff, J., 2017. Early-life inflammation, immune response and ageing. Proceedings of the Royal Society B: Biological Sciences 284, 20170125. https://doi.org/10.1098/rspb.2017.0125

11. Kraaijeveld, A.R., Godfray, H.C.J., 1997. Trade-off between parasitoid resistance and larval competitive ability in *Drosophila melanogaster*. Nature 389, 278–280. https://doi.org/10.1038/38483

12. Kutzer, M. a. M., Kurtz, J., Armitage, S. a. O., 2018. Genotype and diet affect resistance, survival, and fecundity but not fecundity tolerance. Journal of Evolutionary Biology 31, 159–171. https://doi.org/10.1111/jeb.13211

13. Kutzer, M.A.M., Armitage, S.A.O., 2016. The effect of diet and time after bacterial infection on fecundity, resistance, and tolerance in *Drosophila melanogaster*. Ecology and Evolution 6, 4229–4242. https://doi.org/10.1002/ece3.2185

14. Kuznetsova, A., Brockhoff, P.B., Christensen, R.H.B., (2017). lmerTest Package: Tests in 670 Linear Mixed Effects Models. Journal of Statistical Software, 82(13):1–26.

15. Lazzaro, B.P., Sackton, T.B., Clark, A.G., 2006. Genetic Variation in *Drosophila melanogaster* Resistance to Infection: A Comparison Across Bacteria. Genetics 174, 1539– 1554. https://doi.org/10.1534/genetics.105.054593

16. Lawniczak, M.K.N., Barnes, A.I., Linklater, J.R., Boone, J.M., Wigby, S., Chapman, T., 2007. Mating and immunity in invertebrates. Trends in Ecology & Evolution 22, 48–55. https://doi.org/10.1016/j.tree.2006.09.012

17. Lenth, R.V., (2021). emmeans: Estimated Marginal Means, aka Least-Squares 672 Means. R package version 1.6.1.

18. Linder, J.E., Promislow, D.E.L., 2009. Cross-generational fitness effects of infection in *Drosophila melanogaster*. Fly 3, 143–150. https://doi.org/10.4161/fly.8051

19. Lochmiller, R.L., Deerenberg, C., 2000. Trade-offs in evolutionary immunology: just what is the cost of immunity? Oikos 88, 87–98. https://doi.org/10.1034/j.1600-0706.2000.880110.x

20. Mckean, K., Lazzaro, B., 2011. The costs of immunity and the evolution of immunological defense mechanisms. pp. 299–310. https://doi.org/10.1093/acprof:oso/9780199568765.003.0023

21. McKean, K.A., Yourth, C.P., Lazzaro, B.P., Clark, A.G., 2008. The evolutionary costs of immunological maintenance and deployment. BMC Evolutionary Biology 8, 76. https://doi.org/10.1186/1471-2148-8-76

22. Prasad, N.G., Joshi, A., 2003. What have two decades of laboratory life-history evolution studies on *Drosophila melanogaster* taught us? J Genet 82, 45–76. https://doi.org/10.1007/BF02715881

23. R Core Team (2021). R: A language and environment for statistical computing. R Foundation for Statistical Computing, Vienna, Austria. URL: https://www.Rproject.org/.

24. Rion, S., Kawecki, T.J., 2007. Evolutionary biology of starvation resistance: what we have learned from *Drosophila*. Journal of Evolutionary Biology 20, 1655–1664. https://doi.org/10.1111/j.1420-9101.2007.01405.x

25. Rolff, J., Siva-Jothy, M.T., 2003. Invertebrate Ecological Immunology. Science 301, 472–475. https://doi.org/10.1126/science.1080623

26. Rose, M.R., 1984. Laboratory Evolution of Postponed Senescence in Drosophila melanogaster. Evolution 38, 1004–1010. https://doi.org/10.2307/2408434

27. Schmid-Hempel, P., 2005. Evolutionary Ecology of Insect Immune Defenses. Annual Review of Entomology 50, 529–551. https://doi.org/10.1146/annurev.ento.50.071803.130420

28. Schmid-Hempel, P., 2003. Variation in immune defence as a question of evolutionary ecology. Proceedings of the Royal Society of London. Series B: Biological Sciences 270, 357–366. https://doi.org/10.1098/rspb.2002.2265

29. Schulenburg, H., Kurtz, J., Moret, Y., Siva-Jothy, M.T., 2009. Introduction. Ecological immunology. Philosophical Transactions of the Royal Society B: Biological Sciences 364, 3–14. https://doi.org/10.1098/rstb.2008.0249

30. Schwenke, R.A., Lazzaro, B.P., Wolfner, M.F., 2016. Reproduction–Immunity trade-offs in insects. Annual Review of Entomology 61, 239–256. https://doi.org/10.1146/annurev-ento-010715-023924

31. Sheldon, B.C., Verhulst, S., 1996. Ecological immunology: costly parasite defences and trade-offs in evolutionary ecology. Trends in Ecology & Evolution 11, 317–321. https://doi.org/10.1016/0169-5347(96)10039-2

32. Singh, A., Basu, A., Shit, B., Hegde, T., Bansal, N., Prasad, N.G., 2021. Recurrent evolution of cross-resistance in response to selection for improved post-infection survival in *Drosophila melanogaster*. https://doi.org/10.1101/2021.11.26.470139

33. Therneau, T.M., (2020). coxme: Mixed Effects Cox Models. R package version 2.2–668 16.

34. Therneau, T., (2021). A Package for Survival Analysis in R. R package version 3.2–11.

35. Ye, Y.H., Chenoweth, S.F., McGraw, E.A., 2009. Effective but costly, evolved mechanisms of defense against a virulent opportunistic pathogen in *Drosophila melanogaster*. PLOS Pathogens 5, e1000385. https://doi.org/10.1371/journal.ppat.1000385

